# Generation and comparative analysis of full-length transcriptomes in sweetpotato and its putative wild ancestor *I. trifida*

**DOI:** 10.1101/112425

**Authors:** Yonghai Luo, Na Ding, Xuan Shi, Yunxiang Wu, Ruyuan Wang, Lingquan Pei, Ruoyu Xu, Shuo Cheng, Yongyan Lian, Jingyan Gao, Aimin Wang, Jun Tang, Qinghe Cao

**Affiliations:** Plant Functional Genomics, School of Life Sciences, Jiangsu Normal University, Xuzhou City, Jiangsu Province 221116, China; Key Laboratory of Biology and Genetic Improvement of Sweetpotato, Ministry of Agriculture, Xuzhou, Jiangsu 221131, China; Xuzhou Institute of Agricultural Sciences in Jiangsu Xuhuai Region, Xuzhou 221131, Jiangsu, China

## Abstract

Sweetpotato [*Ipomoea batatas* (L.) Lam.] is one of the most important crops in many developing countries and provides a candidate source of bioenergy. However, neither high-quality reference genome nor large-scale full-length cDNA sequences for this outcrossing hexaploid are still lacking, which in turn impedes progress in research studies in sweetpotato functional genomics and molecular breeding. In this study, we apply a combination of second- and third-generation sequencing technologies to sequence full-length transcriptomes in sweetpotato and its putative ancestor *I. trifida*. In total, we obtained 53,861/51,184 high-quality transcripts, which includes 34,963/33,637 putative full-length cDNA sequences, from sweetpotato/*I. trifida*. Amongst, we identified 104,540/94,174 open reading frames, 1476/1475 transcription factors, 25,315/27,090 simple sequence repeats, 417/531 long non-coding RNAs out of the sweetpotato/*I. trifida* dataset. By utilizing public available genomic contigs, we analyzed the gene features (including exon number, exon size, intron number, intron size, exon-intron structure) of 33,119 and 32,793 full-length transcripts in sweetpotato and *I. trifida*, respectively. Furthermore, comparative analysis between our transcript datasets and other large-scale cDNA datasets from different plant species enables us assessing the quality of public datasets, estimating the genetic similarity across relative species, and surveyed the evolutionary pattern of genes. Overall, our study provided fundamental resources of large-scale full-length transcripts in sweetpotato and its putative ancestor, for the first time, and would facilitate structural, functional and comparative genomics studies in this important crop.

## Introduction

The advent of second-generation sequencing (SGS) technologies have revolutionized DNA sequencing and genome/transcriptome studying (Margulies et al., 2005; Shendure and Ji, 2008). In contrast to the first-generation Sanger sequencing, SGS technologies exhibit great advances in reducing cost and exponentially increasing throughput with high sequence accuracy (Sanger et al., 1977; Shendure and Ji, 2008). During the past decades, SGS has become a cost-affordable, routine and widespread method and dramatically accelerate research in biology, especially studies in genomes and transcriptomes. Nevertheless, SGS has disadvantages, one of which is that they generate relatively short reads (i.e., hundreds of base pairs; Koren et al., 2012). In sequencing projects, short reads reduces the accuracy of sequence assembly and make related bioinformatic analyses difficult (Koren et al., 2012). Recently, a third-generation sequencing (TGS) platform [single-molecule real-time (SMRT) sequencer, the PacBio RS; Pacific Biosciences of California, USA] was released (Eid et al., 2009). TGS was designed to overcome the problems of SGS platforms and, most importantly, produced long, albeit relatively low-quality, reads up to 20 kb (Roberts et al., 2013). The long reads of TGS are highly helpful to *de novo* genome and transcriptome assembly in higher organisms and make TGS as an outstanding option in characterizing full-length transcripts (Au et al., 2013; Sharon et al., 2013). The relatively high error rate of TGS might be problematic in sequence alignments and bioinformatics analyses, but could be algorithmically improved and corrected by short and high-accuracy SGS reads (e.g., Koren et al., 2012; Hackl et al., 2014; Li et al., 2014). Therefore, a hybrid sequencing approach combining the SGS and TGS technologies could provide high-quality and more complete assemblies in genome and transcriptome studies, which has been well illustrated previously (e.g., Au et al., 2013; Sharon et al., 2013; Huddleston et al., 2014; Xu et al., 2015).

Long-read or Full-length cDNA sequences are fundamental to studies of structural and functional genomics. First, they provide sequence information of expressed genes and are useful for the functional analyses at the transcriptional and translational levels (Seki et al., 2002). Second, they are useful to identify gene-coding regions within a genome and facilitate to determine the proper orientation, order, and boundary of exons during the predictions of gene models (Imanishi et al., 2004). Third, they are helpful to validate or correct the assembly of a genome and improve gene annotations (Seki et al., 2002; Castelli et al., 2004). Fourth, they are particularly helpful to analyze different transcript isoforms generated by alternate splicing, which is an important process to increase genetic and functional diversity in an organism (e.g., Treutlein et al., 2014). In the past, the generation of full-length cDNA sequences was very expensive, labor intensive, and time consuming because individual cDNA clones should be collected from prepared cDNA libraries and submitted to traditional Sanger sequencing (e.g., Seki et al., 2002; Kikuchi et al., 2003). At present, transcript sequences are most often assembled from high-throughput SGS transcriptome sequencing (i.e., RNA-seq; Mortazavi et al., 2008; Wang et al., 2008), and ideally generated from the SGS/TGS hybrid sequencing approach as described above.

Sweetpotato [*Ipomoea batatas* (L.) Lam.], a Convolvulaceae hexaploid plant, is one of the most important crops in the world because it ensures food supply and safety in many developing countries and provides a candidate source of bioenergy. However, advances in its molecular genetics, genomics, and marker-assisted breeding remain highly restricted due to several reasons. First, sweetpotato is a hexaploid plant with a complex genome (2n = 6x = 90, 3–4 gigabase pairs in genome size; Magoon et al., 1970; Ozias-Akins and Jarret, 1994). By now, there is no high-quality reference genome of sweetpotato available. In 2016, Yang et al announced a preliminary genome assembly for sweetpotato in bioRxiv, but further quality assessment is likely needed (Yang et al. 2016). Second, sweetpotato is a self-incompatible and thus obligate outcrossing species (Martin 1965), resulting in the high heterozygosity of its genome. It is almost impossible to develop typical mapping populations such as F2 and recombinant inbred lines for constructing high-density linkage maps. To date, there is no report in forward genetics (i.e., quantitative trait locus mapping and subsequently map-based cloning) in sweetpotato. Under these situations, RNA-seq (whole transcriptome shotgun sequencing) has been utilized as a promising alternative approach for identifying and analyzing large-scale genes and their expression patterns in sweetpotato (e.g., Wang et al., 2010; Schafleitner et al., 2010; Firon et al., 2013). However, all reported transcriptomic studies in sweetpotato were based on the SGS technologies, and collection and analysis of full-length cDNA sequences remain lacking.

It has long been suggested that sweetpotato was most likely evolved from the diploid ancestor *I. trifida* (Magoon et al., 1970; Ozias-Akins and Jarret, 1994; Srisuwan et al., 2006). Nevertheless, convincing evidence to definitely validate this hypothesis is not provided yet. Since *I. trifida* does not form tuberous roots like sweetpotato does, comparative analysis between sweetpotato and *I. trifida* may provide important insights into the mechanism of tuberization, as well as the evolution or domestication of this important crop. Although a preliminary genome assembly of *I. trifida* has been released in 2015 (Hirakawa et al. 2015), large-scale collection and analysis of its full-length cDNA sequences are not reported.

In this study, we constructed full-length cDNA libraries from sweetpotato and *I. trifida*, and performed SMRT and SGS sequencing to generate large-scale full-length transcripts for either species. Comprehensive intraspecific and interspecific analyses were carried out, which has provided valuable resources for genomics studies in sweetpotato and its putative ancestor *I. trifida*.

## Results and Discussion

### SMRT- and Illumina-based RNA sequencing and error correction

To obtain a representative full-length transcriptome for sweetpotato and *I. trifida*, respectively, single molecule real-time (SMRT) sequencing was performed using a Pacific RSII sequencing platform. For either organism, different tissues were collected and mixed from a single plant and used for mRNA extractions. Three size-fractionated, full-length cDNA libraries (1-2kb, 2-3kb, and >3kb) were constructed and subsequently sequenced in four SMRT cells (1, 2, and 1 cell, respectively; Fig. 1a,b). In sweetpotato, we obtained 220,035 reads of insert (total bases: 701,923,565), which included 49.9% of full-length non-chimeric and 46.6% of non-full-length reads (Fig. 1c and Table 1); while in *I. trifida*, 195,188 reads of insert (total bases: 527,497,043), 52.1% and 43.9% of which are full-length non-chimeric and non-full-length reads, respectively (Fig. 1d and Table 1).

**Figure 1.**
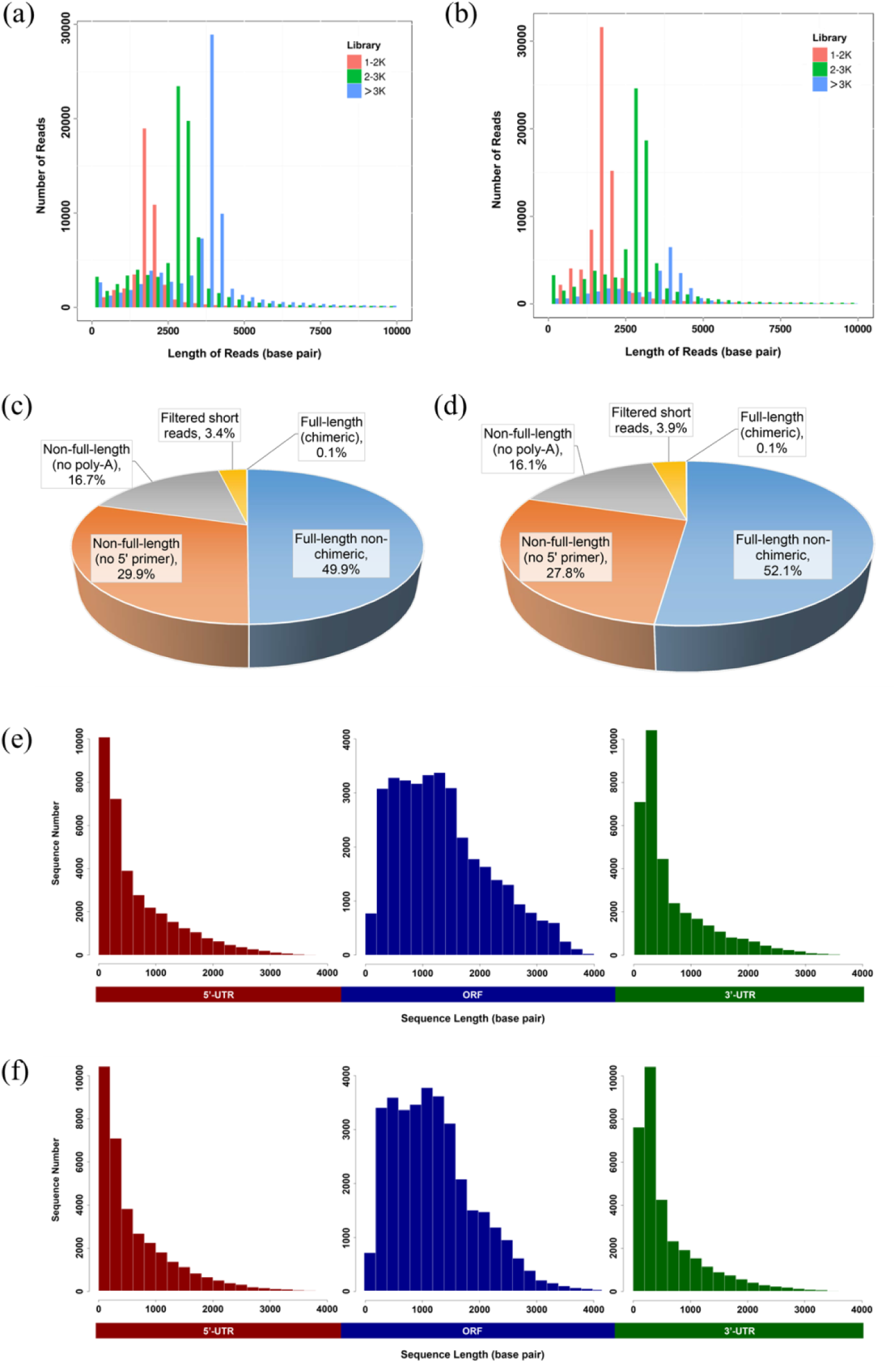
Summary of PacBio RS II single-molecule real-time (SMRT) sequencing. Number and length distributions of 220,035 reads in sweetpotato (**a**) and 195,188 reads in *I. trifida* (**b**) from different PacBio libraries (fractionated size: 1-2, 2-3, >3kb); Proportion of different types of PacBio reads in sweetpotato (**c**) and *I. trifida* (**d**); Number and length distributions of characteristic components of 34, 963 non-redundant full-length transcripts in sweetpotato (**e**) and 33,637 in *I. trifida* (**f**), UTR, untranslated regions; ORF, open reading frame.

**Table 1.** Summary of PacBio- and Illumina-based RNA sequencing in this study.

|  | *I. batatas* | *I. trifida* |
| --- | --- | --- |
| Reads of insert of PacBio sequencing | 220,035 | 195,188 |
| Bases of insert of PacBio sequencing (bp) | 701,923,565 | 527,497,043 |
| Reads of Illumina sequencing for correction | 71,360,785 | 39,372,131 |
| Bases of Illumina sequencing for correction (bp) | 17,972,706,252 | 11,772,267,169 |
| Number of non-full-length PacBio reads | 102,510 | 85,680 |
| Number of full-length non-chimeric PacBio reads | 109,814 | 101,630 |
| Average length of full-length non-chimeric PacBio reads (bp) | 8,641 | 8,488 |
| Number of non-redundant transcripts after correction | 53,861 | 51,184 |
| N50 of non-redundant transcripts after correction (bp) | 2,933 | 2,642 |
| Mean of non-redundant transcripts after correction (bp) | 2,421 | 2,190 |
| Number of non-redundant full-length transcripts after correction | 34,963 | 33,637 |

As SMRT sequencing generates a high error rate, it is necessary to perform error correction, which includes self-correction by iterative clustering of circular-consensus reads and correction with high-quality SGS short reads. To this end, cDNA libraries were prepared from the same samples used for SMRT sequencing and deep RNA sequencings were conducted using an Illumina Hiseq2500 platform. In total, 71,360,785 and 39,372,131 clean reads (total bases: 17,972,706,252 and 11,772,267,169, respectively) were obtained and used to correct the SMRT reads in sweetpotato and *I. trifida*, respectively (Table 1). After error correction, redundant transcripts were removed. Finally, we obtained 53,861 non-redundant transcripts for sweetpotato (named as Ib53861; N50: 2,933bp; mean: 2,421bp) and 51,184 for *I. trifida* (named as It51184; N50: 2,642bp; mean: 2,190bp; Supplementary Fig. S1).

### Predictions of open reading frames (ORFs), simple sequence repeats (SSRs), and long non-coding RNAs (lncRNAs)

In total, 104,540 and 94,174 ORFs were predicted out of Ib53861 and It51184, respectively, and their length distributions were analyzed (Supplementary Fig. S2). Those transcripts containing complete coding sequences (CDSs) as well as 5’-and 3’-UTR (untranslated regions) were defined as full-length transcripts. Respectively, 34,963 and 33,637 full-length transcripts were identified for *I. batatas* (named as Ib34963) and *I. trifida* (named as It33637), and the number and length distributions of their characteristic components were investigated (Fig.1e,f).

SSR markers represent one of the most widely used molecular markers in numerous organisms. Here we detected 25,315 and 27,090 SSRs out of Ib53861 and It51184, respectively, and found that most of them were with mono-, di-, or tri-nucleotide repeats (Table 2; Supplementary Table S1,S2). Considering the high assembly quality of SMRT-derived sequences, the here detected SSRs would be useful for marker-assisted breeding and genetic analysis in sweetpotato and *I. thifida*.

**Table 2.** Prediction of simple sequence repeats (SSRs) out of our transcript datasets.

|  | **Ib53861** | **It51184** |
| --- | --- | --- |
| Total number of sequences examined | 53,841 | 51,164 |
| Total size of examined sequences (bp) | 130,402,110 | 112,068,291 |
| Total number of identified SSRs | 25,319 | 27,090 |
| Number of SSR containing sequences | 17,340 | 17,127 |
| Number of sequences containing more than 1 SSR | 5,401 | 5,448 |
| Number of SSRs present in compound formation | 2,542 | 4,700 |
| Mono- nucleotide | 11,640 | 14,031 |
| Di- nucleotide | 6,942 | 6,571 |
| Tri- nucleotide | 6,092 | 5,784 |
| Tetra- nucleotide | 427 | 432 |
| Penta- nucleotide | 124 | 154 |
| Hexa- nucleotide | 94 | 118 |

LncRNAs are an emerging hot topic in biology and has been found to be functional as key regulators in a wide spectrum of biological processes. In this study, we identified 417 and 531 lncRNAs in sweetpotato and *I. trifida*, respectively (Supplementary Table S3,S4), which provides candidate lncRNAs for further functional characterizations in this worldwide crop.

### Functional annotation of our transcripts and sorting of transcription factors

Transcripts in Ib53861 and It51184 were functionally annotated and classified according to sequence similarities using BLASTx or tBLASTx (E-value ⩽1e-5) against different protein and nucleotide databases (Supplementary Table S5). Overall, 97.25% of Ib53861 and 97.34% of It51184 transcripts were successfully annotated in considered databases, and the successful rates in a single database ranged from 41.67% to 96.46% (Table 3). These results indicate that most of the genes in our datasets are truly transcribed sequences and likely functional genes in sweetpotato and/or *I. trifida*.

**Table 3.**
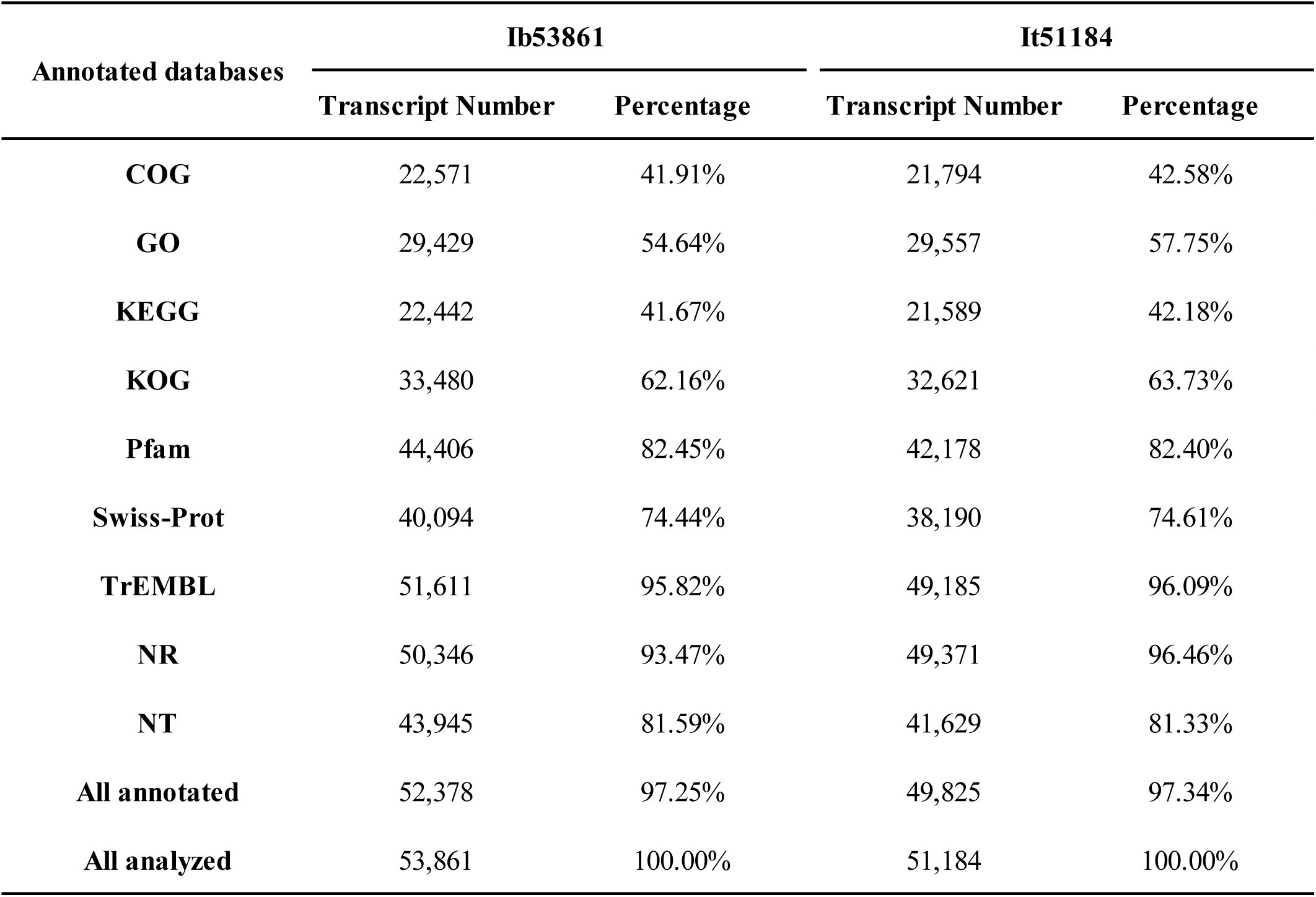
Annotation of our transcript datasets to public databases. COG, clusters of orthologous groups; GO, gene ontology; KEGG, kyoto encyclopedia of genes and genomes; Pfam, a large collection of protein families; Swiss-Prot, a well-annotated and manually checked protein database; TrEMBL, an automatically annotated protein database; NR, a NCBI non-redundant protein database; NT, a NCBI nucleotide database.

| Annotated databases | Ib53861 |  | It51184 |  |
| --- | --- | --- | --- | --- |
|  | Transcript Number | Percentage | Transcript Number | Percentage |
| **COG** | 22,571 | 41.91% | 21,794 | 42.58% |
| **GO** | 29,429 | 54.64% | 29,557 | 57.75% |
| **KEGG** | 22,442 | 41.67% | 21,589 | 42.18% |
| **KOG** | 33,480 | 62.16% | 32,621 | 63.73% |
| **Pfam** | 44,406 | 82.45% | 42,178 | 82.40% |
| **Swiss-Prot** | 40,094 | 74.44% | 38,190 | 74.61% |
| **TrEMBL** | 51,611 | 95.82% | 49,185 | 96.09% |
| **NR** | 50,346 | 93.47% | 49,371 | 96.46% |
| **NT** | 43,945 | 81.59% | 41,629 | 81.33% |
| **All annotated** | 52,378 | 97.25% | 49,825 | 97.34% |
| **All analyzed** | 53,861 | 100.00% | 51,184 | 100.00% |

Transcription factors (TFs) are key components involved in the transcriptional regulatory system in an organism and thus hot targets in various research themes. To highlight TFs available in our transcript datasets, we examined 52 TF gene families and identified 1464 and 1475 putative TF members out of Ib53861 and It51184, respectively (Supplementary Table S6,S7). When compared our TF numbers to those in several other plants, we found that they were likely underrepresented, to some extent, for most TF gene families (Supplementary Table S7). This is not abnormal since the expression of most of TFs are spatiotemporally controlled and thus hard to be detected simultaneously in a transcriptional level.

### Comparison of our transcripts with datasets in other plants

To assess the similarity of our transcripts with genes of other plants, we compared Ib53861 and It51184 to several released CDS datasets derived from corresponding genome sequencing projects, including *I. batatas* (Yang et al., 2016), *I. trifida* (Hirakawa et al., 2015), *Ipomoea nil* (Hoshino et al., 2016), *Nicotiana tabacum* (Sierro et al., 2013), *Solanum tuberosum* (The Potato Genome Sequencing Consortium, 2011), *Glycine max* (Schmutz et al., 2010), *Arabidopsis thaliana* (The Arabidopsis Genome Initiative, 2000), and *Oryza sativa* (International Rice Genome Sequencing Project, 2005). As shown in Figure 2, our analysis has revealed several implications. First, most of transcripts in Ib53861 and It51184 were homologous in whatever cutoff values (low, medium, or high). This suggests that two species share a largely same gene reservoir and thus are phylogenetically closed. Second, we found remarkably less homologs when used the Yang’s CDS dataset, in contrast to the Yang’s genomic dataset, as a subject database (Fig. 2). This indicates that the assembly quality of Yang’s contigs seem significantly higher than their CDS predictions. Third, high proportions of transcripts could find their homologous sequences in a low cutoff value, but not in a medium or high cutoff, across the here investigated organisms including rice, a monocot plant. This result indicates that (i) a large number of genes in these species still share short motifs despite of their phylogenetically far distance, although great sequence divergences have apparently occurred during evolution; (ii) nevertheless, there are still some genes showing evolutionarily high conservation across lineages. Fourth, the descending order of similarity of large-scale CDS sequences from other species to sweetpotato was *I. trifida, I. nil, N. tabacum, S. tuberosum, G. max, A. thaliana, and O. sativa*. Notably, the cDNA dataset of *I. nil* looks highly homologous to that of sweetpotato or *I. trifida*.

**Figure 2.**
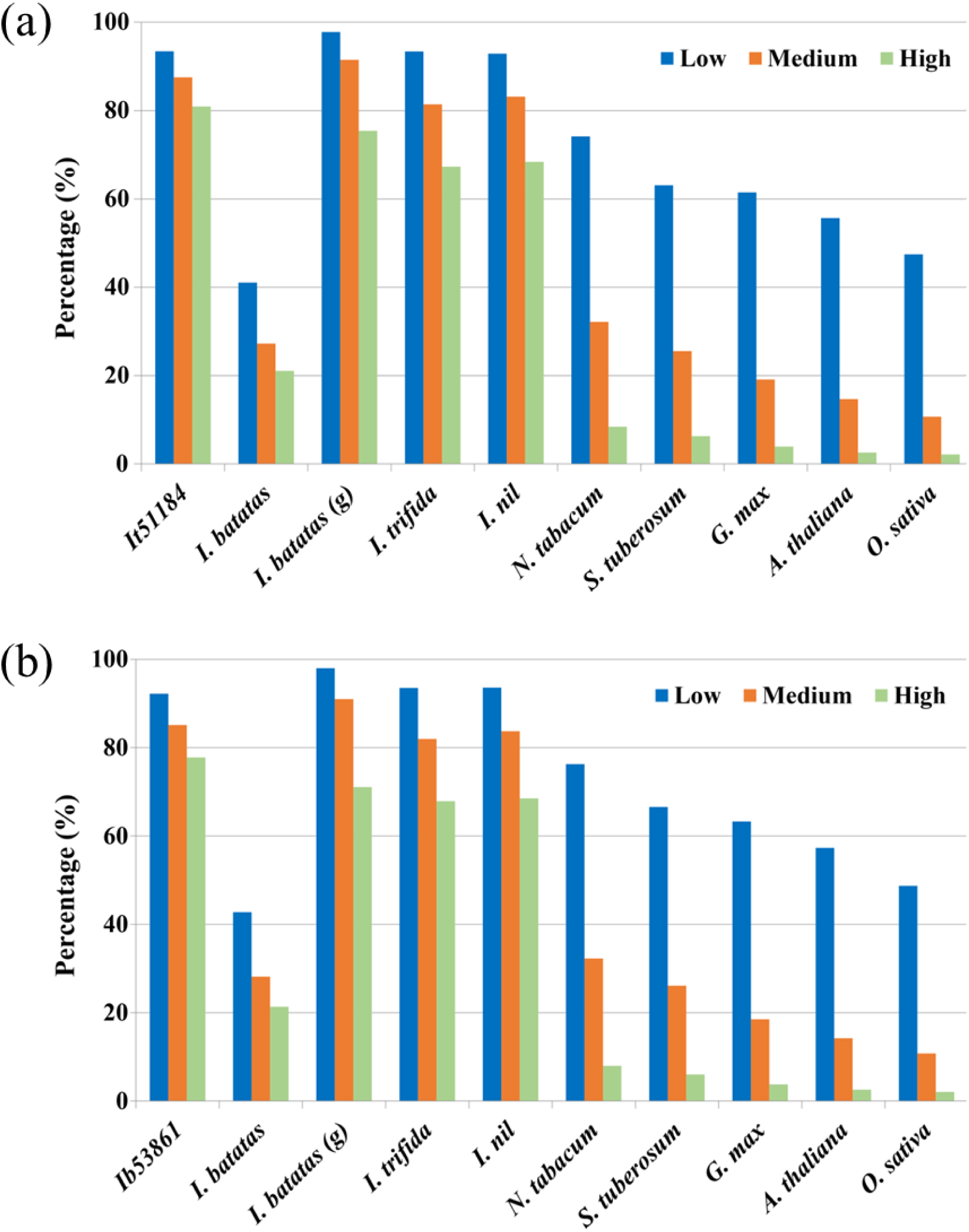
Comparison of our transcripts with genes of other plants. BLASTx or tBLASTx searches were performed and percentages of successful hits under different criteria were shown for Ib53861 (**a**) and It51184 (**b**). All subject datasets were predicted coding sequences in the indicated species except *I. batatas* (*g*), which is the dataset of Yang’s sweetpotato genomic scaffolds. Low means low similarity (number of identical matches >= 20 and percentage of identical matches >= 60%), while Medium and High refer to 60/80% and 100/90%, respectively.

### Analysis of exon–intron structures of full-length transcripts

Long-read or full-length cDNA sequences are useful for structural genomics studies, such as analysis of exon-intron structures. Using a stringent cutoff, we mapped each of our full-length transcripts (i.e., Ib34963 and It33637) onto a best-matched contig in the corresponding species and analyzed its possible exon-intron structure. To examine whether our BLASTn-based pipeline could work well, we firstly analyzed the exon-intron structures of genes in *A. thaliana*, one of the best well-characterized organism. As shown in Figure 3, our results (i.e., genes with introns, exons per gene, mean exon or intron size, and pattern of distribution of exon or intron size) are similar to those reported (The Arabidopsis Genome Initiative, 2000). We further manually and randomly examined exon-intron structures of 50 genes with various intron numbers in our analysis and compared to those shown at TAIR (http://www.arabidopsis.org). Only a very few differences in exon-intron predictions were found, which could be mainly attributed to differences in the release versions of genome and filtering parameters. These data validated our working pipeline.

**Figure 3.**
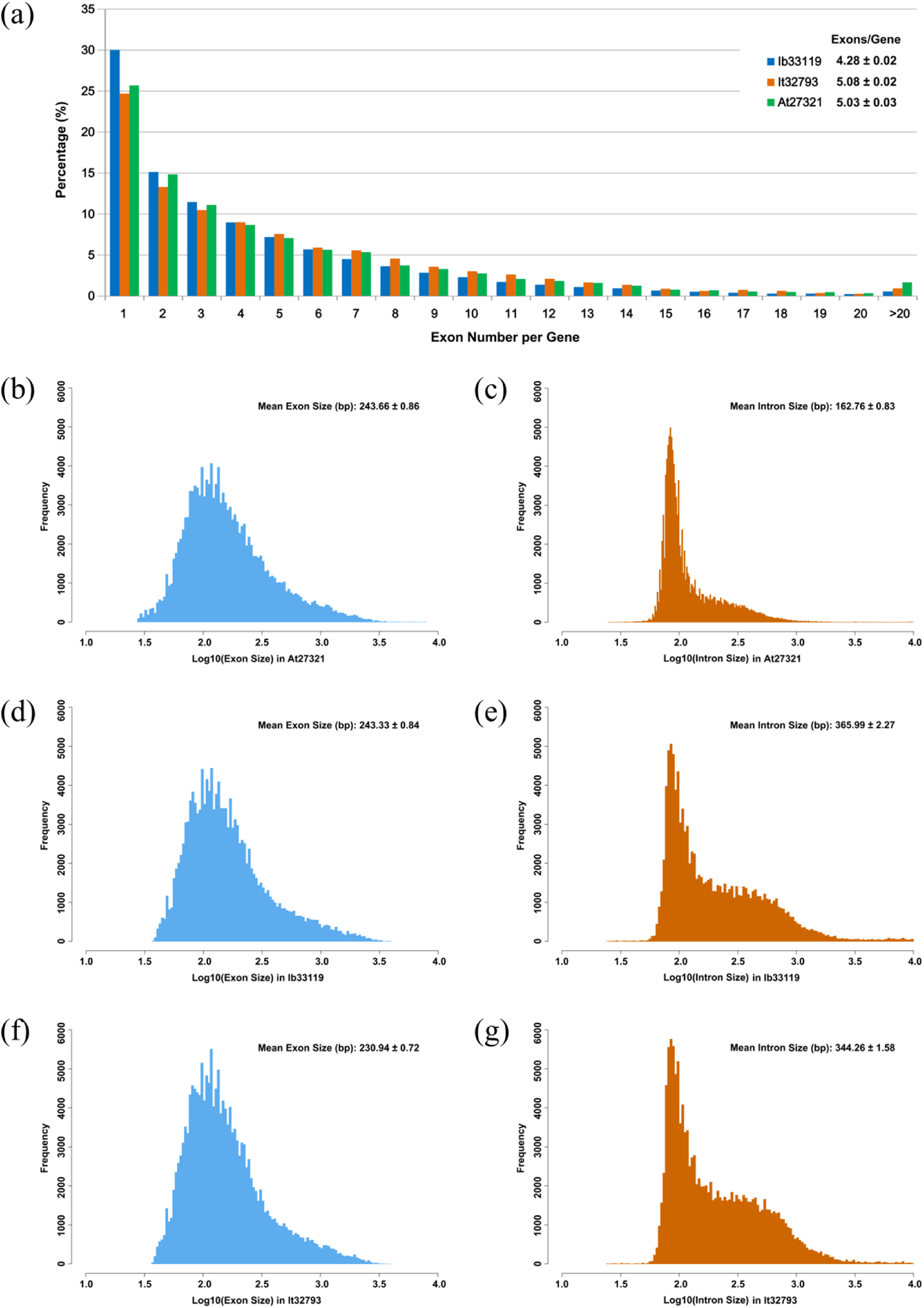
Analysis of exon-intron structures in *A. thaliana*, sweetpotato and *I. trifida*. (**a**) Percentage distribution of genes with different exon number. Distribution of exon size and intron size in different species, (**b**,**c**) *A. thaliana*; (**d**,**e**) sweetpotato; (**f**,**g**) *I. trifida*.

In total, 33,119 (95%) and 32,793 (97.5%) full-length transcripts in sweetpotato and *I. trifida*, respectively, were successfully analyzed, and detailed information of exon-intron structures was provided (Supplementary Table S8,S9). The average exon number per gene in sweetpotato was 4.28, which was significantly smaller than 5.08 in *I. trifida*, and 5.03 in *A. thaliana* (either P < 0.0001), although their distributions were comparable across three species (Fig. 3b). The average exon size in sweetpotato was 243.66bp, which was close to 243.33bp in *A. thaliana* (P = 1.000) and significantly bigger to 230.94bp in *I. trifida* (P < 0.0001), and their distributions were only slightly different (Fig. 3c,e,g). The means of intron sizes were significantly different: 365.99bp in sweetpotato, 344.26bp in *I. trifida*, and 162.76bp in *A. thaliana*, respectively (all P < 0.0001); and the distribution pattern of sweetpotato was similar to that of *I. trifida*, but apparently distinguishable from that of *A. thaliana* (Fig. 3d,f,h).

We are aware that some of the predicted exon-intron structures may be imprecise or even erroneous because of the relative low quality of sweetpotato scaffolds or *I. trifida* contigs. However, for the first time, our analyses provide the preliminary information of exon-intron structures in large-scale full-length cDNA sequences in sweetpotato, which is useful for genomics and comparative genomics studies (as examples shown in Supplementary Fig. S3).

### Estimation of Ka/Ks ratios in putative orthologous gene pairs between sweetpotato and ***I. trifida***

It was thought that the diploid plant *I. trifida* was the ancestor of the autohexaploid crop sweetpotato. To search candidate genes involved in the evolution or domestication of sweetpotato, we investigated the selective pattern of genes between sweetpotato and *I. trifida*. We compared the full-length transcript datasets of Ib34963 and It33637, removed those transcripts with multiple possible orthologs, and finally identified 1269 putative orthologous gene pairs between sweetpotato and *I. trifida*. Subsequently, the nonsynonymous and synonymous substitution rates (Ka and Ks, respectively) as well as the Ka/Ks ratio of each gene pair were calculated (Supplementary Table S10). As shown in Figure 4, the Ka/Ks values of most of the gene pairs were smaller than one, and only 56 gene pairs exhibited Ka/Ks > 1 and 9 of them had Ka/Ks > 2. These data suggest that most genes are likely under purifying selection during the evolution or domestication of sweetpotato. The roles of those genes under strong positive selection must await for further study.

**Figure 4.**
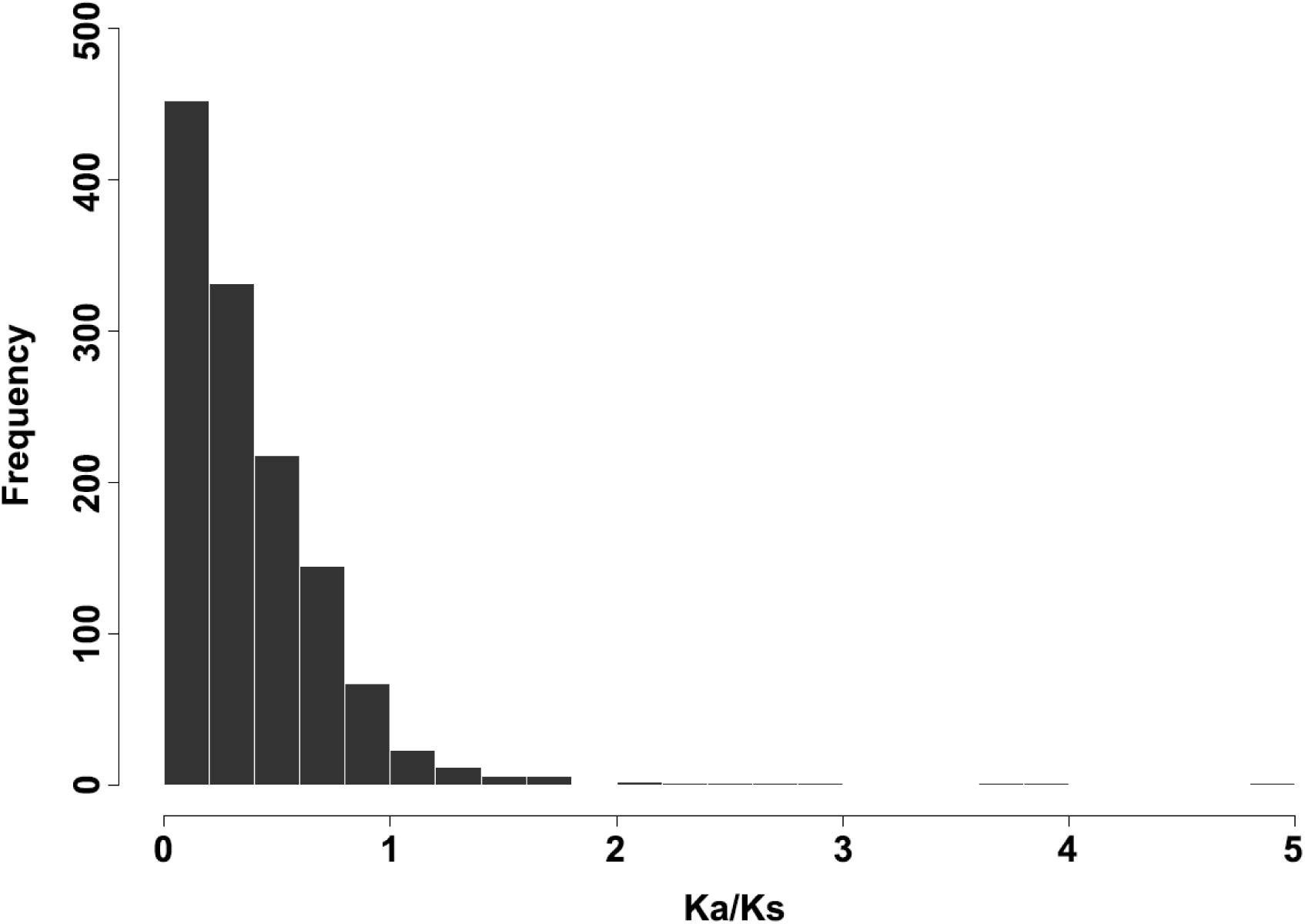
Distribution of Ka/Ks ratios in 1269 putative orthologous gene pairs between sweetpotato and *I. trifida*.

## Conclusion

Full-length cDNA sequences are fundamental resources to studies of structural, functional, and comparative genomics but hardly available in sweetpotato and its relatives. In this study, we sequenced full-length transcriptomes in both sweetpotato and its putative ancestor *I. trifida*, by using a combination of PacBio SMRT sequencing and Illumina RNA sequencing technologies. In total, 53,861 transcripts (including 34,963 putative full-length ones) and 51184 transcripts (including 33,637 putative full-length ones) were produced in sweetpotato and *I. trifida*, respectively, Predictions of ORFs, SSRs, lncRNAs, transcription factors, gene structures, and functional annotations as well as comparative analyses across plant species were performed. Our analyses have provided a comprehensive transcriptome database for either sweetpotato or *I. trifida*, which would facilitate structural, functional and comparative genomics studies in this worldwide crop.

## Methods and Materials

### Plant material and RNA preparation

*Xushu18*, one of the most widely cultivated sweetpotato varieties in China, was selected for RNA sequencing. Tissues of young leaves, mature leaves, shoots, stems, fibrous roots, initiating tuberous roots, expanding tuberous roots, and mature tuberous roots from one individual were collected and pooled together in approximately equivalent weights. Comparably, tissues of young leaves, mature leaves, shoots, stems, and roots of a diploid *I. trifida* plant (provided by the Chinese Sweetpotato Research Institute, Xuzhou, China) were collected and pooled. Collected samples were frozen in liquid nitrogen immediately after collection and stored at −80°C before use.

Total RNAs were extracted using Tiangen RNA preparation kits (Tiangen Biotech, Beijing, China) following the provided protocol. RNA quality and quantity were determined using a Nanodrop ND-1000 spectrophotometer (NanoDrop Technologies, Wilmington, DE, USA) and a 2100 Bioanalyzer (Agilent Technologies, Palo Alto, CA, USA). Qualified RNA samples were used for constructing cDNA libraries subsequently.

### PacBio cDNA library construction and third-generation sequencing

cDNA was synthesized using the SMARTer PCR cDNA Synthesis Kit, optimized for preparing full-length cDNA (Takara Clontech Biotech, Dalian, China). Size fractionation and selection (1-2kb, 2-3kb, and >3kb) were performed using the BluePippin^™^ Size Selection System (Sage Science, Beverly, MA). The SMRT bell libraries were constructed with the Pacific Biosciences DNA Template Prep Kit 2.0. SMRT sequencing was then performed on the Pacific Bioscience RS II platform using the provided protocol.

### Illumina cDNA library construction and second-generation sequencing

The SGS cDNA libraries were constructed using a NEBNext® Ultra^™^ RNA Library Prep Kit for Illumina® (NEB, Beverly, MA, USA), following the manufacturer’s protocol. Qualified libraries were applied to NGS using an Illumina Hiseq 2500 (Illumina, San Diego, CA, USA) to generate 125bp paired-end sequence reads (2 × 125bp). High-throughput sequencing (both TGS and SGS) reported in this study was performed in the Biomarker Technology Co. (Beijing, China).

### Quality filtering and error correction of PacBio long reads

The TGS subreads were filtered using the standard protocols in the SMRT Analysis software suite (http://www.pacificbiosciences.com), and reads of insert (ROIs) were obtained. After examining for poly(A) signal and 5’ and 3’ adaptors, full-length and non-full-length cDNA reads were recognized. Consensus isoforms were identified using the algorithm of iterative clustering for error correction and further polished to obtain high-quality consensus isoforms. The raw Illumina SGS reads were filtered to remove adaptor sequences, ambiguous reads with ‘N’ bases, and low-quality reads. Afterward, error correction of low-quality isoforms was conducted using the SGS reads with the software proovread 2.13.841. Redundant isoforms were then removed to generate a high-quality transcript dataset for each species (i.e., Ib53861 for *I. batatas* and It51184 for *I. trifida*, respectively) using the program CD-HIT 4.6.142 (http://weizhongli-lab.org/cd-hit/).

### Functional annotation of transcripts

Functional annotations were conducted by using BLASTX (cutoff E-value ≤ 1e-5) against different protein and nucleotide databases of COG (Clusters of Orthologous Groups), GO (Gene Ontology), KEGG (Kyoto Encyclopedia of Genes and Genomes), Pfam (a database of conserved Protein families or domains), Swiss-prot (a manually annotated, non-redundant protein database), TrEMBL (an automatically annotated protein database), NR (NCBI non-redundant proteins), and NT (the NCBI nucleotide database). For each transcript in each database searching, the functional information of the best matched sequence was assigned to the query transcript.

### Predictions of ORF, SSR, and long non-coding RNAs

To predict the open reading frames (ORFs) in transcripts, we used the package TransDecoder v2.0.1 (https://transdecoder.github.io/) to define putative coding sequences (CDS). The predicted CDS were searched and confirmed by BLASTX (E-value ⩽1e-5) against the protein databases of NR, SWISS-PROT, and KEGG. Those transcripts containing complete ORFs as well as 5’-and 3’-UTR (untranslated regions) were designated as full-length transcripts. To identify putative simple sequence repeats (SSRs) in our assembled sequences, the tool MISA (MIcroSAtellite identification tool; http://pgrc.ipk-gatersleben.de/misa) was employed. Only transcripts with ⩾500 bp in size were used in SSR detection. To predict long non-coding RNAs (lncRNAs), we used PLEK (predictor of long non-coding RNAs and messenger RNAs based on an improved k-mer scheme; https://sourceforge.net/projects/plek).

### Identification of transcription factors and in comparison to those in other plants

For a transcription factor gene family, the Hidden Markov Model (HMM) profile of Pfam domain (if available) was downloaded from the Pfam database (http://pfam.xfam.org; Finn et al., 2016) and applied as a query to survey all predicted proteins out of our transcript datasets using HMMER (http://www.hmmer.org). All identified non-redundant transcripts with expected values less than 1.0 were collected and confirmed the existence of featured domains by searching the NCBI Conserved Domain database (https://www.ncbi.nlm.nih.gov/Structure/cdd/wrpsb.cgi). If there were no HMM profile available for a gene family, all protein sequences belonged to the gene family in *A. thaliana* were downloaded (http://www.arabidopsis.org) and used as query sequences to search our predicted protein datasets using BLASTp (E-value ⩽1e-10). Subsequently, the identified transcripts were confirmed in the NCBI Conserved Domain database. The confirmed transcripts of each gene family were compiled and the number was compared with those in several other plant species (the data were retrieved from the Plant Transcription Factor Database v4.0 (PlantTFDB): http://planttfdb.cbi.pku.edu.cn; Jin et al., 2017). For *I. nil*, the sorting of transcription factors was done as same as that for sweetpotato since there are no related data available in PlantTFDB yet.

### Analysis of sequence similarity to genes in other plants

To survey the sequence similarity to genes in other plant species, we compared our transcripts of Ib53861 and It51184, respectively, to the predicted cDNA or protein datasets in *I. batatas* (http://public-genomes-ngs.molgen.mpg.de/SweetPotato; Yang et al., 2016), *I. trifida* (http://sweetpotato-garden.kazusa.or.jp; Hirakawa et al., 2015), *I. nil* (https://www.ncbi.nlm.nih.gov/genome/46552; Hoshino et al., 2016), *N. tabacum* (https://www.ncbi.nlm.nih.gov/genome/425; Sierro et al., 2013), *S. tuberosum* (http://solanaceae.plantbiology.msu.edu; The Potato Genome Sequencing Consortium, 2011), *G. max* (https://phytozome.jgi.doe.gov; Schmutz et al., 2010), *A. thaliana* (http://www.arabidopsis.org; The Arabidopsis Genome Iniative, 2000), and *O. sativa* (http://rice.plantbiology.msu.edu; International Rice Genome Sequencing Project, 2005). BLASTx or tBLASTx (E-value ⩽1e-5) was used for the homologous search and the best-matched sequence was picked up for each query transcript. The results were further filtered with different parameters: low similarity (number of identical matches >= 20 and percentage of identical matches >= 60%), while medium or high similarity refers to 60/80% or 100/90%, respectively.

### Analysis of exon–intron structures of full-length transcripts

The full-length transcripts in Ib34259 and It33637 were mapped to the genomic contigs in *I. batatas* and *I. trifida*, respectively, using BLASTn. The alignment hits between a given transcript and the best-matched contig were collected and filtered firstly using the parameter: number of identical matches >= 25 and percentage of identical matches >= 90%. The mismatched or redundant alignment hits were further filtered and the exon-intron structure for each transcript were predicted using in-house scripts. Afterward, the numbers and lengths of exons and introns were statistically analyzed in a transcriptome level. To improve and validate our in-house scripts, we downloaded the datasets of coding and genomic sequences of *A. thaliana* and adapted our working pipeline in this well-characterized species.

### Estimation of Ka/Ks in putative orthologous gene pairs

The OrthoMCL algorithm with default settings was applied to identify orthologous gene pairs between Ib34259 and It33637 (http://orthomcl.org; Li et al., 2003; Fischer et al., 2011). A transcript in one species with multiple possible orthologues in the other species was excluded. For each orthologous gene pair, sequence alignment were done with the MUSCLE program (http://drive5.com/muscle; Edgar 2004). After that, the nonsynonymous and synonymous substitution rates (Ka and Ks, respectively) were estimated using the YN00 program of Phylogenetic Analysis by Maximum Likelihood (PAML) based on the basic model (Yang 1997).

### Availability of sequence data

The PacBio SMRT reads and the Illumina SGS reads generated in this study have been submitted to the BioProject database of National Center for Biotechnology Information (accession number XXXXXXX).

## Acknowledgements

This study was jointly supported by the Priority Academic Program Development of Jiangsu Higher Education Institutions, National sweet potato industry and research system (CARS-11-B-02), Natural Science Foundation of Jiangsu Province (BK20141146), National Key Laboratory of Plant Molecular Genetics (Y409Z111U1), and Education Department of Jiangsu Province (KYLX15_1468).

---

**Supplementary Figure S1.**
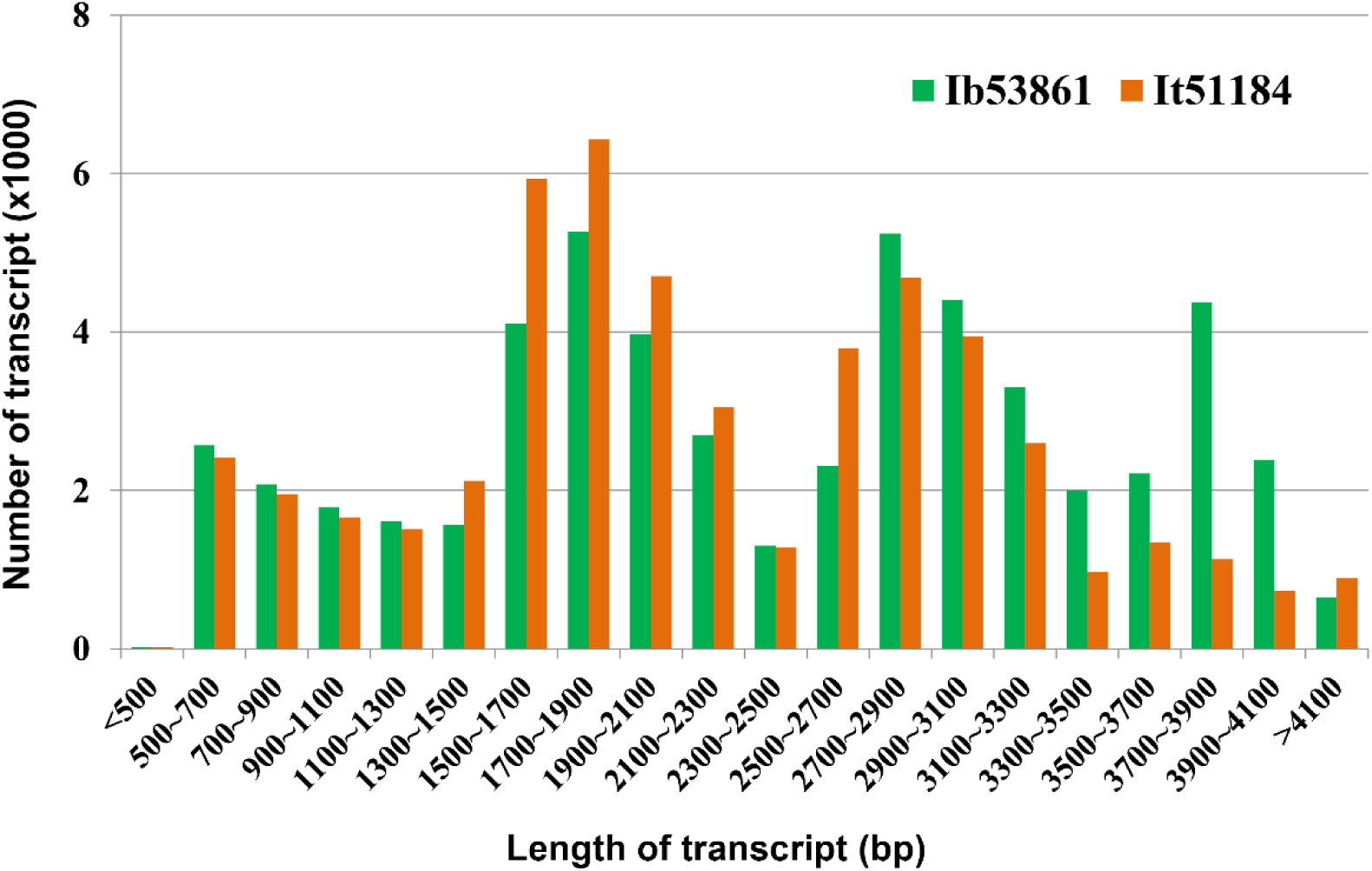
Number and length distributions of 53,861 non-redundant transcripts in sweetpotato (Ib53861) and 51,184 in *I. trifida* (It51184).

**Supplementary Figure S2.**
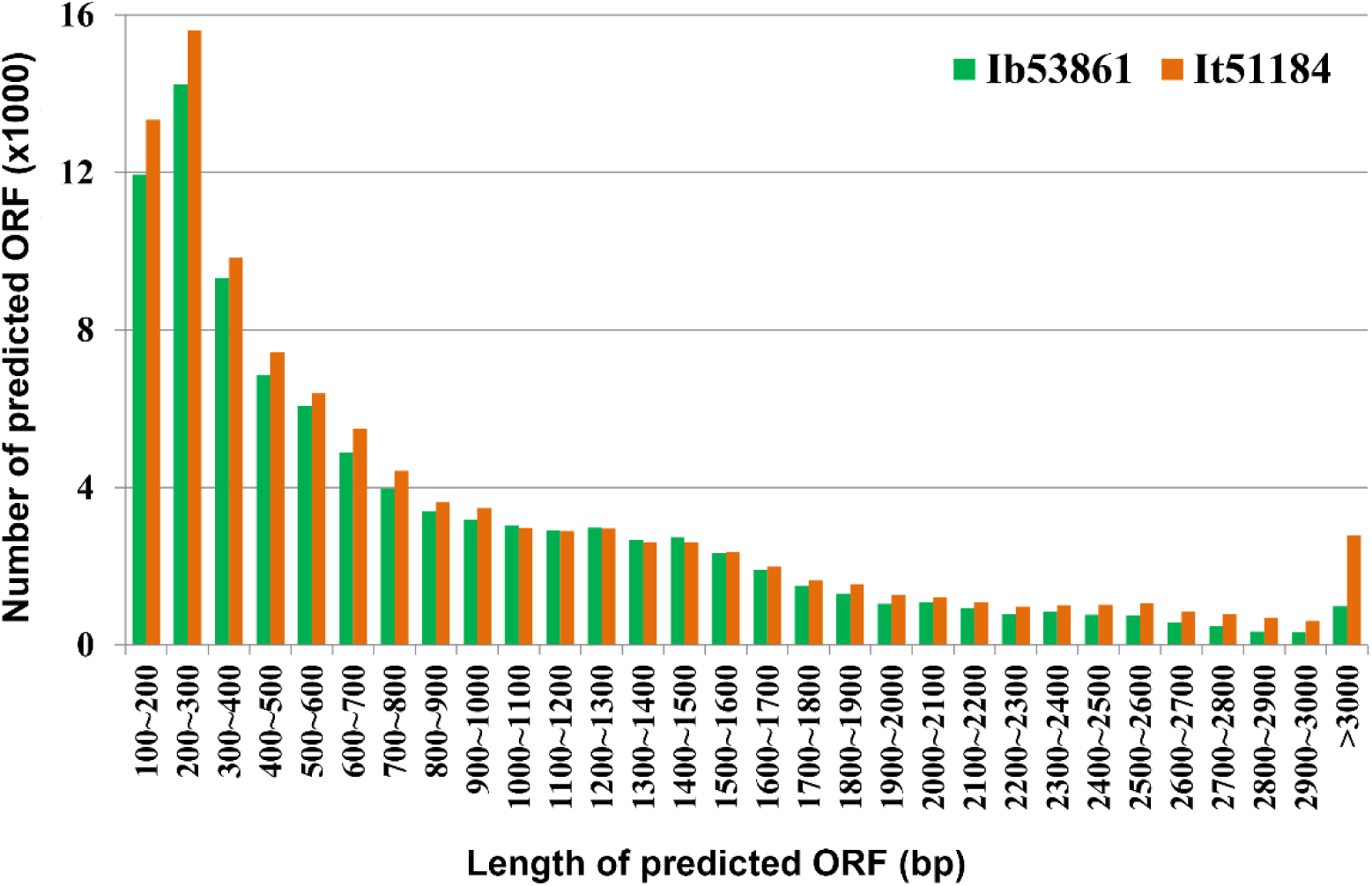
Number and length distributions of predicted open reading frames (ORFs) in 53,861 non-redundant transcripts in sweetpotato (Ib53861) and 51,184 in *I. trifida* (It51184).

**Supplementary Figure S3.**
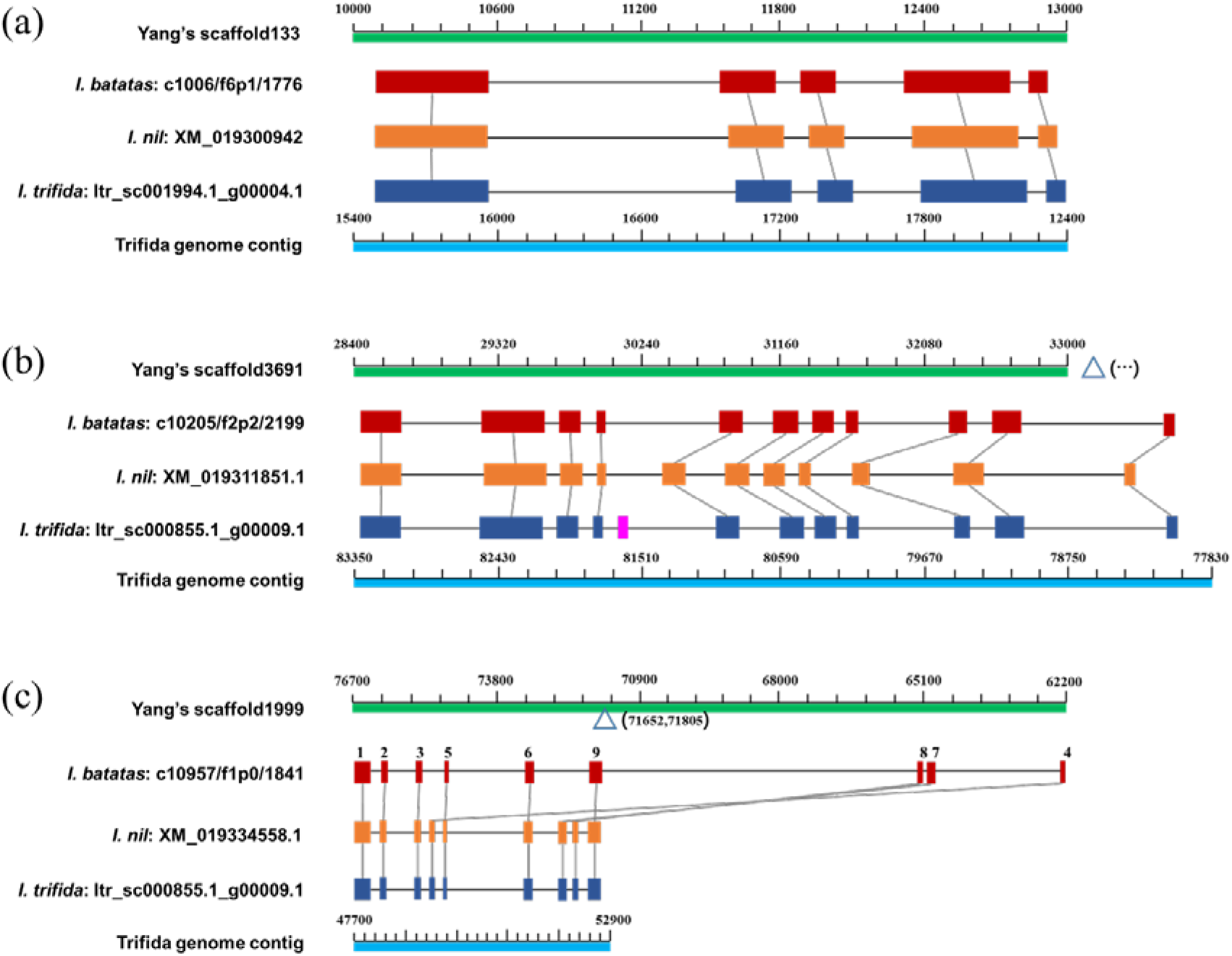
Examples of comparative analyses of genes across sweetpotato, *I. nil*, and *I. trifida*. (**a**) An example showing highly conserved gene models; (**b**) An example showing exon variation (i.e., one additional exon exists in I. trifida); (**c**) An example showing a likely abnormal gene model in sweetpotato, which could be due to assembly error of genomic sequences.

